# 17α Estradiol promotes plasticity of spared inputs in the adult amblyopic visual cortex

**DOI:** 10.1101/773374

**Authors:** Deepali C. Sengupta, Crystal L. Lantz, M.A. Karim Rumi, Elizabeth M. Quinlan

**Affiliations:** Neuroscience and Cognitive Science Program, University of Maryland, College Park, MD 20742; Department of Biology, University of Maryland, College Park, MD 20742; Department of Pathology and Laboratory Medicine, University of Kansas Medical Center, Kansas City, KS 66160; Institute for Reproduction and Perinatal Research, University of Kansas Medical Center, Kansas City, KS 66160

## Abstract

The promotion of structural and functional plasticity by estrogens is a promising therapy to enhance central nervous system function in the aged. However, how the sensitivity to estrogens is regulated across brain regions, age and experience is poorly understood. To ask if estradiol treatment impacts structural and functional plasticity in sensory cortices, we examined the acute effect of 17α Estradiol in adult Long Evans (LE) rats following chronic monocular deprivation, a manipulation that reduces the strength and selectivity of deprived eye vision. Chronic monocular deprivation decreased thalamic input from the deprived eye to the binocular visual cortex and accelerated short-term depression of the deprived eye pathway, without changing the total density of excitatory synapses. Importantly, we found that the classical estrogen receptors ERα and ERβ are robustly expressed in the adult visual cortex, and that a single dose of 17α Estradiol increased the size of excitatory postsynaptic densities, reduced the expression of parvalbumin and decreased the integrity of the extracellular matrix. Furthermore, 17α Estradiol enhanced experience-dependent plasticity in the amblyopic visual cortex, and promoted response potentiation of the pathway served by the non-deprived eye. The promotion of plasticity at synapses serving the non-deprived eye may reflect selectivity for synapses with an initially low probability of neurotransmitter release, and may inform applications to remap spared inputs around a scotoma or a cortical infarct

## Introduction

Age-related constraints in synaptic plasticity have long been associated with a decrease in circulating sex hormones. Accordingly, estrogen therapy (estradiol valerate) is reported to prevent age-related cognitive impairment and improve verbal memory in oophorectomized premenopausal women^1^. Similar estrogen therapy reduces the development of all-cause dementia if initiated by age 65. The benefits of estrogens have also been demonstrated in animal models. In aged rhesus macaques, estrogen replacement (estradiol cypionate) following ovariectomy (OVX) reverses age-related cognitive impairment^2^. In OVX rats, treatment with17α-estradiol (αE2), a stereoisomer of the main circulating estradiol (E2), improved object/place recognition and enhanced spatial working memory^3^. E2 treatment also improved water maze memory retrieval and reduced the severity score after traumatic brain injury in adult OVX mice (11-13 months^4^). The majority of previous work was performed on gonadectomized subjects, to control for circulating hormone levels, and estrogen treatment in intact subjects produces more modest effects^5, 6^. Importantly, gonadally-intact non-human primates experience less multi-synaptic bouton loss with age in the prefrontal cortex than OVX cohorts^7^, suggesting that surgically induced menopause may have more severe neurological consequences than natural aging.

Circulating estrogens likely promote the maintenance of synapse number and neuronal excitability that is permissive for activity-dependent synaptic plasticity. Consequently, an age-related decrease in circulating E2 correlates with a decrease in synaptic glutamate receptor density, and inhibition of E2 biosynthesis induces acute spine synapse loss^8, 9^. Similarly, dendritic spine density and the magnitude of long term potentiation in the hippocampus and cortex are increased by E2 in intact and OVX adult rats^10,11,12^. Importantly, E2 delivery to intact male or intact and OVX female rats increases paired pulse depression, enhances synaptic strength, and promotes activity-dependent synaptic potentiation at excitatory synapses^13,14,15^.

The impact of estrogens on central nervous system function has been elucidated primarily in hippocampus and prefrontal cortex, areas known to express estrogen receptors (ERs) in young (3-4 months), middle-aged (9-11 months), and aged (19-24 months) female rats^16, 17, 18^. Classical estrogen signaling is mediated by cytoplasmic receptors that translocate to the nucleus to bind DNA and regulate transcription. However, E2 also enhances cognition and plasticity independently of transcription^19, 20^. ERs associated with the plasma membrane and/or cytoplasmic organelles likely mediate the non-transcriptional regulation of synaptic function^21, 22^. The canonical estrogen receptors, estrogen receptor α (ERα) and estrogen receptor β (ERβ) are present at multiple sites in neurons^16, 23, 24^. ERα has been localized to nuclei, cytoplasm and the presynaptic compartment of pyramidal neurons in hippocampal CA1, and immunogold electron microscopy (EM) localizes ERα to dendritic spines^25^. Immunogold EM also localizes ERβ to pre- and postsynaptic compartments of asymmetric synapses in CA1 in female rats, which persist following OVX and are increased by E2 treatment^17^. ERα and ERβ puncta co- localize with the postsynaptic scaffold protein PSD95, and E2 delivery recruits PSD95 to synapses in cultured neurons^26^. ERα is also present in approximately one-third of the inhibitory boutons that innervate pyramidal neuron somata in hippocampal CA1^27^. Presynaptic ERα co-localizes with cholecystokinin, is associated with clusters of synaptic vesicles, and is mobilized toward the synapse by E2. Importantly, expression of membrane ERs persists in the adult hippocampus and prefrontal cortex following OVX^16, 17, 18^. In contrast, there is little consensus on the distribution or role of ERs in primary sensory cortices of adults, due in part to conflicts between previous reports, which also differ in age, species, sex and gonadal state^28, 29, 30, 31, 32^.

Here we examine the distribution of estrogen receptors in V1 of adult male and female LE rats following chronic monocular deprivation (cMD). cMD initiated at eye opening mimics the presence of a unilateral congenital cataract from birth and induces severe amblyopia that is highly resistant to reversal in adulthood^33, 34^. Importantly, in the binocular region of the amblyopic cortex, synapses serving the deprived eye are weak, sparse and untuned, while synapses serving the non-deprived eye retain normal strength, density and stimulus selectivity^35^. In addition to inducing deprivation amblyopia, the reduction in synaptic density and cortical function mimics changes observed during normal aging or in response to ischemic damage. We asked if a acute treatment with a single low dose of estradiol can enhance plasticity in V1 of adult amblyopic rodents. We employ αE2, which has 100-fold lower efficacy for nuclear ERs of both subtypes and a preferential affinity for ERα^36^ and has been shown to induce ~2 fold more synaptogenesis in the hippocampus than an equivalent dose of E2^37, 38^.

## Materials and Methods

### Subjects

Long Evans (LE) Rats (strain 006, RRID:RGD_2308852) were purchased from Charles River Laboratories (Raleigh, NC). Equal numbers of adult (>postnatal day 180, >P180) males and females were used. Animals were raised in 12/12 hour light/dark cycle and experiments were performed, or subjects were sacrificed, 6 hours into the light phase. Brains from 8-12 week old female ERβ^−/−^ rats were provided by M.A. Karim Rumi of the University of Kansas Medical Center^39^. All procedures were approved by the University of Maryland Institutional Animal Care and Use Committee and were carried out in accordance with the Guide for the Care and Use of Laboratory Animals.

### Monocular deprivation

P14 LE rat pups were anesthetized with ketamine/xylazine (100 mg/10 mg/kg, intraperitoneal). The margins of the upper and lower lids of one eye were trimmed and sutured together using a 5-0 suture kit with polyglycolic acid (CP Medical). Subjects were returned to their home cage after recovery at 37°C for 1-2 hours and disqualified in the event of suture opening.

### Antibodies

The following antibodies/dilutions were used: rabbit anti-estrogen receptor α (ERα; ThermoFisher, RRID: AB_325813, 1:1000); mouse anti-estrogen receptor β (ERβ; ThermoFisher; AB_2717280, 1:1000); mouse anti-parvalbumin (PV; Millipore; RRID:AB_2174013, 1:1000); mouse anti-Post Synaptic Density 95kd (PSD95; ThermoFisher; RRID: AB_325399, 1:200); rabbit anti-phospho- Serine 831-GluR1 (pS831; Sigma Aldrich; RRID:AB_1977218, 1:1000); followed by appropriate secondary antibodies: goat anti-rabbit and anti-mouse IgG conjugated to Alexa-488 or 647 (Life Technologies; RRID:AB_143165, RRID:AB_2535805, 1:300). The specificity of the ERα antibody was validated by the vendor through immunolabeling in positive (ERα^+^/ERβ^+^ T547D^36,40^) and negative cells (ERα^−^/ERβ^−^ Hs578T^41^). We performed our own confirmation of the specificity of the ERβ, see results.

### Reagents

HSV-H129 EGFP (strain 772; University of Pittsburgh Center for Neuroanatomy and Neurotropic Viruses (CNNV)) and HSV-H129 mCherry (strain 373; CNNV) were diluted 1:1 with diH_2_O and 3μl was injected intraocularly (right eye: mCherry; left eye: EGFP) for anterograde delivery to primary visual cortex. 4’,6-Diamidine-2’-phenylindole dihydrochloride (DAPI, Sigma; 1:10000) was used to visualize cell nuclei. Chondroitin sulfate proteoglycans (CSPGs) were visualized with fluorescein wisteria floribunda agglutinin (WFA, Vector Labs; 1:1000). 17α Estradiol (αE2; Sigma Aldrich) was diluted to 15 μg/kg in sesame oil (veh., Sigma Aldrich) and administered subcutaneously (s.c.) to awake animals.

### Immunohistochemistry

Subjects were perfused transcardially with phosphate buffered saline (PBS) followed by 4% paraformaldehyde (PFA) in PBS. The brain was post-fixed in 4% PFA for 24 hours followed by 30% sucrose for 48 hours and cryo-protectant solution (0.58 M sucrose, 30% (v/v) ethylene glycol, 3 mM sodium azide, 0.64 M sodium phosphate, pH 7.4) for 24 hours prior to sectioning. Coronal sections (40 μm) were made on a Leica freezing microtome (Model SM 2000R). Sections (ML: 4mm AP: −6.72 mm DV: 1.5mm) were blocked with 4% normal goat serum (NGS) in PBS for 1 hour. Antibodies were presented in blocking solution for 24 hours, followed by appropriate secondary antibodies for 2 hours. Immunolabeling for ERα and ERβ was absent when antibodies were pre- absorbed with antigen (*not shown*).

### Confocal imaging and analysis

Images were acquired on a Zeiss LSM 710 confocal microscope. Tiled photomontages of ERα, ERβ, and ERα / ERβ with DAPI were constructed with MosaicJ (FIJI, NIH) from individual images (2.8 mm^2^) acquired at 5x magnification (Zeiss Plan-neofluar 5x/0.16, NA=0.16). Co-localization of ERα and ERβ with DAPI was analyzed in single z-section images (70.86 × 70.86 μm 550 μm depth from cortical surface (corresponding to layer 4 of V1), 3 coronal sections / subject, 1 region of interest (ROI) / hemisphere) taken at 40X (Zeiss Plan-neofluar 40x/1.3 Oil DIC, NA=1.3), using the JACoP plugin in FIJI (NIH). Pearson’s correlation coefficient was used to calculate the covariance of two fluorescent signals independently of fluorescence intensity. HSV anterograde viral tracer signal was visualized in single z-section images (2.8 mm^2^ or 1.4 mm^2^) acquired at 5x or 10x magnification (Zeiss Plan-neofluar 10x/0.30, NA=0.30) and a mean intensity profile was calculated using FIJI. The cortical distribution of WFA and PV immunoreactivity was determined in a z-stack (9 × 7.5 μm sections, 3 coronal sections / subject, 1 ROI/hemisphere) acquired at 10X. Maximal intensity projections (MIPs; 500 μm width, 900 μm depth from cortical surface) were used to obtain mean intensity profiles in FIJI. For PSD95 and pS831 staining, MIPs of z-stacks (40 slices × 0.9 μm images; 3 coronal sections / subject, 1 ROI/hemisphere) were acquired at 100X (Zeiss Plan-neofluar 100x/1.3 Oil DIC, NA=1.3). PSD95 and pS831 puncta were selected using size exclusion parameters defined by unbiased quantification for each marker following the construction of a cumulative distribution of puncta size, and setting a 10% lower bound and 90% upper bound. In our acquisition setup, resolution (lambda*N.A.) = 200 nm, and our upper bound of 0.6 microns^2^ is ~ 3*resolution. The lower limit is imposed to exclude subresolution, single pixels/stochastic noise from the analysis. Puncta were identified in MIPs (28.34 × 28.34 × 40 μm z-stack images, 550 μm depth from cortical surface) based on fluorescence thresholding (autothreshold) in FIJI.

### *Acute* In Vivo *Recordings*

Visually evoked potentials (VEPs) were recorded from the binocular region of primary visual cortex (V1b) of adult rats contralateral to the chronically deprived eye. Rats were anesthetized with 3% isoflurane in 100% O_2_ and a 3 mm craniotomy was produced over V1b (centered 3 mm medial from midline and 7 mm posterior from Bregma). A 1.8 mm 16-channel platinum-iridium linear electrode array (~112 μm site spacing, 250 kΩ) was inserted perpendicular to V1b (dorsal/ventral: 1.8 mm). Recordings under 2.5% isoflurane in 100% O_2_ commenced 30 minutes after electrode insertion. Local field potentials were acquired via a RZ5 amplifier (Tucker Davis Technology) with a 300 Hz low pass filter and a 60 Hz notch filter. VEPs were evoked through passive viewing of 100 × 1 second trials of square-wave gratings (0.05 cycles per degree (cpd), 100% contrast, reversing at 1 Hz, via MATLAB (Mathworks) with Psychtoolbox extensions. Average VEP waveforms were calculated for 100 stimulus presentations and were assigned to layers based on waveform shape. VEP amplitude was measured from trough to peak in MATLAB^42^. To examine the short-term plasticity of VEP amplitude, each eye was individually presented with full field flashes (90 cd/m^2^), alternating with 0 cd/m^2^ every 0.5 seconds. Single trial VEP responses were normalized to the first evoked response. To examine the response of the amblyopic cortex to repetitive patterned visual stimulation, subjects received passive binocular stimulation of 200 phase reversals of 0.05 cycles per degree (cpd), 100% contrast gratings, 45 degrees, reversing at 1 Hz. After 24 hours, VEPs were evoked from each eye (originally deprived and non-deprived eye separately), in response to the familiar (45 degrees) and a novel (135 degrees) grating stimulus.

### Experimental Design and Statistical Analysis

Primary visual cortex was defined with anatomical landmarks (dimensions of the dorsal hippocampal commissure, deep cerebral white matter tract, and the forceps major of the corpus callosum). Modest shrinkage due to fixation and cryoprotection reduced vertical depth to 900 μm^35^. Fluorescent puncta were identified using size exclusion parameters defined by unbiased quantification for each marker following the construction of a cumulative distribution of puncta size, and setting a 10% lower bound and 90% upper bound. An unpaired two-tailed Student’s T-test was used to determine the significance between two independent experimental groups, and a paired Student’s T-test was used for two measurements within the same subject. One-way ANOVA was used to determine significance between three independent groups. Repeated measures ANOVA, with between group comparisons, was used to determine the significance of more than two measures within the same subjects, followed by a Tukey-Kramer honestly significant difference *post hoc* for pairwise comparisons if p < 0.05 (JASP). A Kolmogorov-Smirnov test (K-S Test) was used to determine the significance between the distributions of two independent data sets. Multi-dimensional K-S Test was used for distributions with two independent measurements (MATLAB). Statistical significance (p<0.05) is represented as asterisks in figures, and data is presented as mean ± standard error (mean±SEM). Where statistical significance was observed (p<0.05), the effect size was calculated (Cohen’s d) as mean of group 1 mean – group 2 mean/combined standard deviation of groups 1 and 2, with d < 0.2 considered a small effect, d >0.2 - <0.5 considered medium and d>0.8 considered a large effect.

## Results

### Effects of cMD on thalamocortical innervation of V1

Estradiol (E2) treatment has been shown to promote short-term synaptic plasticity, increase the number and size of excitatory synapses, and lower the threshold for activity-dependent potentiation^14, 37, 43^. We therefore first asked how each of these potential targets of E2 is impacted in an animal model in which amblyopia is induced by chronic monocular deprivation (cMD) from eye opening to adulthood (P14 - >P180). Neurons in the binocular region of primary visual cortex (V1b) in rodents have a preference for stimulation from the contralateral eye, as approximately twice as many thalamocortical afferents serve the contralateral as ipsilateral eye^44^. A representative example of eye-specific innervation from the thalamus to the cortex, revealed by dual intraocular injection of the anterograde trans-neuronal label HSV-H129, confirms > 1.5 fold innervation of layer 4 from the contralateral than the ipsilateral eye in adult binocular controls (Average Fluorescence ±SEM; Ipsi HSV-EGFP 54.61±0.22, Contra HSV-mCherry 81.33±0.85; Fig.1B). Brief monocular deprivation shifts ocular preference away from the deprived eye, due in part to a reduction in the number of thalamic afferents serving the deprived eye innervating V1b^45, 46^. Accordingly, cMD significantly decreases the thalamocortical innervation from the deprived contralateral eye, reducing the initial contralateral bias (average Fluorescence ±SEM; Ipsi HSV-EGFP 40.26±0.25, Contra HSV- mCherry 47.38±0.22, Fig.1B). cMD also induced an expansion of the cortical territory innervated by the non-deprived eye into the monocular region of V1 (V1m), as previously observed in felines^47^.

**Figure 1.**
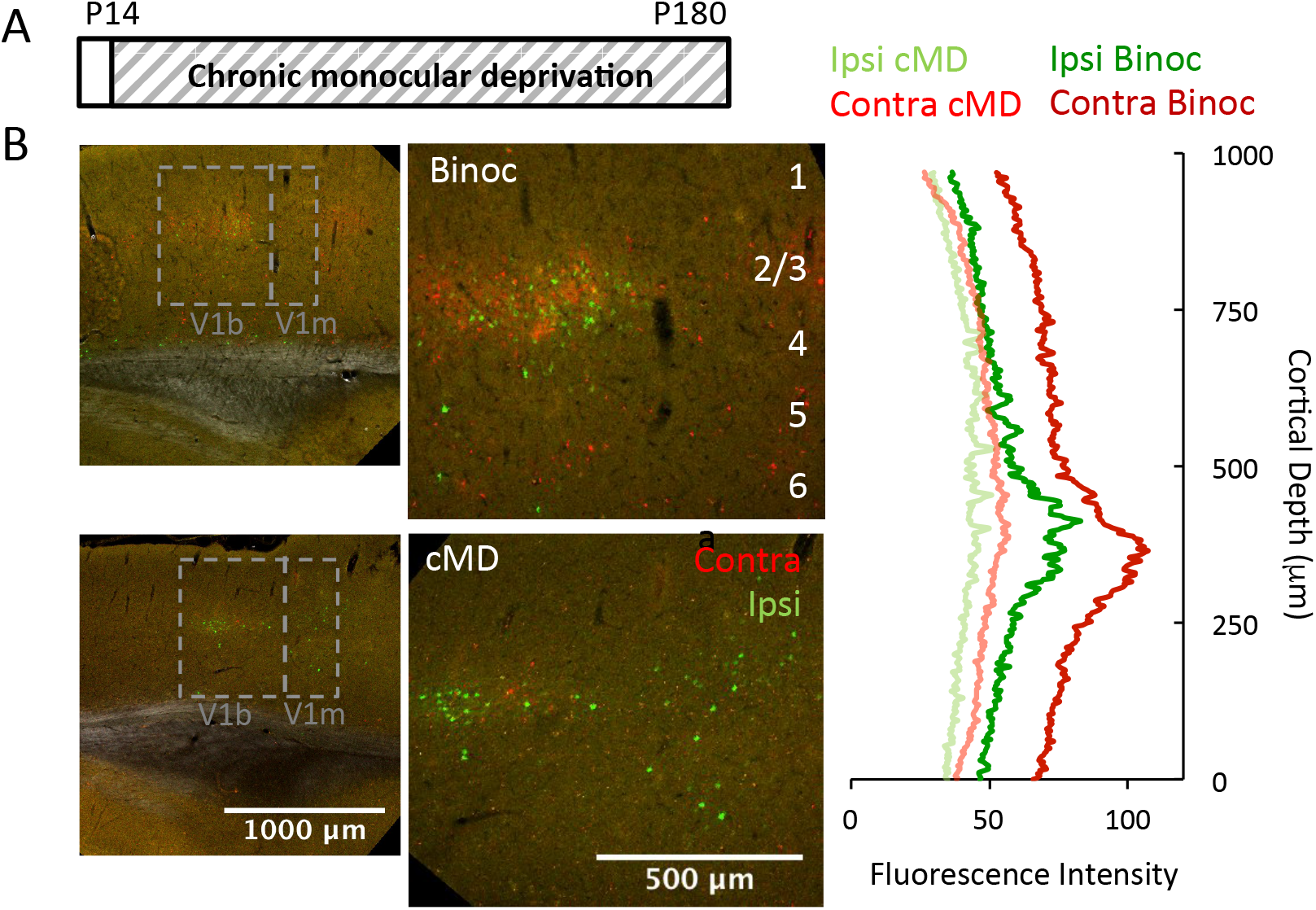
Chronic monocular deprivation decreases thalamic input to primary visual cortex. A. Experimental timeline. Subjects received monocular deprivation from eye opening (~P14) to adulthood (>P180). B. Left: Representative confocal double fluorescent micrographs of primary visual cortex (V1) following intravitreal delivery of H129 373, mCherry into right/contralateral and H129 772, EGFP into left/ipsilateral eye of a control binocular (top) and a cMD subject (bottom). V1 location (here and all other figures) ML: 4mm AP: −6.72 mm DV: 1.5mm; ROI: 2000μm × 2000μm; left: 5x mag with 0.6x digital zoom; right: 900μm × 1000μm; 10x mag with 0.6x digital zoom). Right: Quantification of mCherry and EGFP fluorescence in V1b (900X100 ROI, presented by vertical depth) of higher magnification images shows reduction in trans-neuronal label following cMD.

### Quantitative immunofluorescence of PSD95 and pS831 following cMD

To ask if cMD impacted the number and/or size of excitatory synapses in V1b, we quantified the intensity and distribution of the scaffold protein PSD95. PSD95 is detected in nascent synaptic connections, and plays a fundamental role in many aspects of excitatory synapse maturation and LTP induction/expression by co-localizing critical synaptic effectors. Quantitative immunofluorescence revealed that cMD significantly reduced PSD95 puncta size distribution in deprived and non-deprived primary visual cortex (AVG±SEM; Binoc 0.092±0.01 μm^2^ vs. Contra cMD 0.087±0.01 μm^2^, p<0.001, K-S Test, Cohen’s *d* =0.51; Binoc 0.092±0.01 μm^2^ vs. Ipsi cMD 0.088±0.01 μm^2^; p<0.001, K-S Test, Cohen’s *d* = 0.39, Fig.1C; males versus females: F(1,16) = 0.29, p = 0.87, 2-way ANOVA). However, we observed no difference in PSD95 puncta number in V1b contralateral or ipsilateral to the chronically-deprived eye (AVG±SEM; Binocular Control (Binoc) 333.36±44.19, Contralateral (Contra cMD) 320.65±54.46, Ipsilateral (Ipsi cMD) 297.87±53.71; males versus females: F(1,16) = 0.76, p = 0.14, 2-way ANOVA; n=8, Fig.2A).

**Figure 2.**
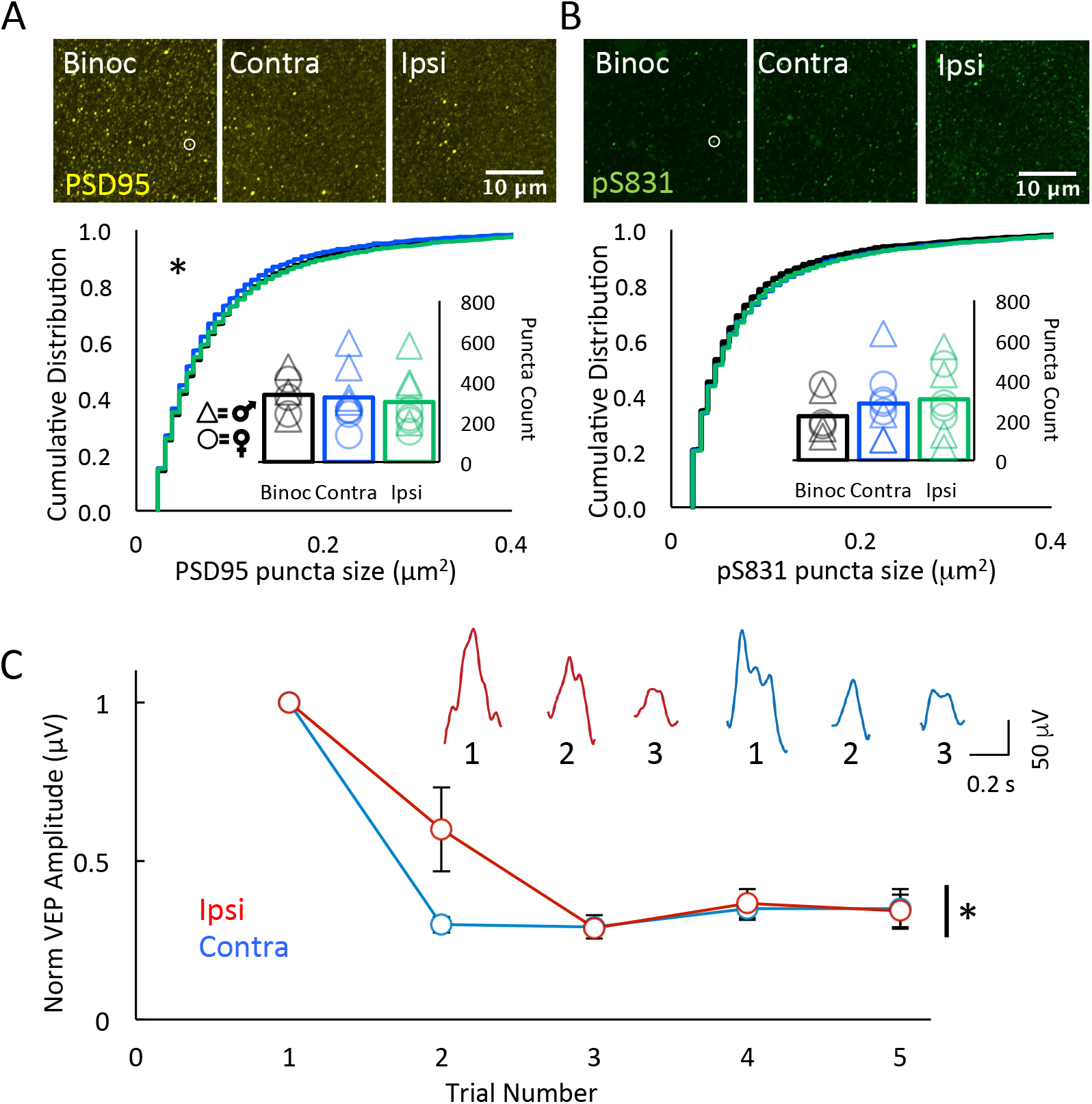
Chronic monocular deprivation decrease the size of PSD95 puncta and regulates the response to repetitive visual stimulation. A. Top: Fluorescent micrographs of PSD95 (yellow, representative puncta in white circle) in V1b of binocular control (Binoc, left) and following cMD (Contra cMD, middle; Ipsi cMD, right). ROI 500 μm from surface; 28.34 μm × 28.34 μm × 40 μm, 100x mag with 3x digital zoom; maximal intensity projection (MIP; 40 × 1μm z-steps). Bottom: Significant decrease in PSD95 immunoreactive puncta size *=p<0.001, K-S Test Contra and Ipsi relative to Binoc. No change in puncta number (inset) in V1b following cMDed. Males (triangles) vs. females (circles): 2-way ANOVAs size: F(1,16) = 0.29, p = 0.87; number: F(1,16) = 0.76, p = 0.14, 2-way ANOVA; Binoc n = 3 males, 3 females, cMD n = 4 males, 4 females. B. Top: Fluorescent micrographs of pS831 (green, representative puncta in white circle) in V1b of binocular control (Binoc, left) and following cMD (Contra cMD, middle; Ipsi cMD, right). ROI as in 2A. Bottom: No change in pS831 immunoreactive puncta size or number (inset) in V1b contralateral or ipsilateral to the cMDed eye. Males (triangles) vs. females (circles): 2-way ANOVA Size: F(1,16) = 0.17, p = 0.69; Number: F(1,16) = 0.78, p = 0.39,)F(1,16) = 0.29, p = 0.87; Binoc n = 3 males, 3 females, cMD n = 4 males, 4 females. C. VEP amplitudes acquired from layer 2/3 in response to single full field light flash to ipsilateral (non-deprived, red) and contralateral eye (deprived, blue) normalized to amplitude of first response. Contralateral eye VEPs depress more rapidly than ipsilateral eye VEPs. Inset: representative raw, single VEP waveforms in response to 3 consecutive light flashes (full field, 0.5 second 90 cd/m^2^, 0.5 second 0 cd/m^2^) to ipsilateral (red) and contralateral (blue) eyes. Second VEP response normalized to first, AVG±SEM; Ipsi: 0.59±0.13, Contra: 0.29±0.02; Repeated measures ANOVA, *p < 0.001, F = 47.683, between groups, p = 0.007, F = 15.773, n = 4 subjects.

To ask if cMD impacted the activity of signaling pathways known to regulate glutamate receptor function, we examined the intensity and distribution of the GluA1 subunit of the α-amino-3-hydroxy- 5-methyl-4-isoxazolepropionic acid receptor subtype of glutamate receptor (AMPAR) phosphorylated on residue Serine 831 (pS831). Ca^2+^/calmodulin-dependent protein kinase II (CaMKII) and protein kinase C (PKC)-dependent phosphorylation of Serine 831 of the GluA1 subunit increases single channel conductance and promotes LTP^48^. Importantly, rapid and transient phosphorylation of S831 is induced *in vivo* by salient stimulation^49, 50^. However, following cMD, we observed no difference in the size or number of pS831 puncta in V1b contralateral or ipsilateral to the occluded eye (AVG±SEM; Size: Binoc 0.078±0.003 μm^2^, Contra cMD 0.078±0.006 μm^2^, Ipsi cMD 0.077±0.006 μm^2^. Males versus females: F(1,16) = 0.17, p = 0.69, 2-way ANOVA; Number: Binoc 227.82±48.40, Contra cMD 283.68±64.10, Ipsi cMD 305.01±66.20. Males versus females: F(1,16) = 0.78, p = 0.39, 2-way ANOVA; Binoc n=6, cMD n = 8, Fig.2B). Thus, the deprivation-induced reduction in thalamic input to cortex does not reduce excitatory synaptic density, likely reflecting an increase in the number of other classes of excitatory synapses.

### VEP responses following cMD

To ask if cMD impacted short-term activity-dependent synaptic plasticity, we examined changes in the amplitude of visually evoked local field potentials (VEPs) in response to repetitive visual stimulation of each eye. Microelectrode array recordings of the VEP isolated from layer 2/3 of V1b contralateral to the occluded eye (deprived hemisphere) were acquired in response to repetitive flash stimuli (full field, 0.5 second 90 cd/m^2^, 0.5 second 0 cd/m^2^) and amplitudes were normalized to the first VEP. Full field flashes of light were used to maximally stimulate synapses in the amblyopic cortex, as in prior work chronic monocular deprivation severely compromises spatial acuity ^33, 35, 51^. The shape of the VEP waveform is a more reliable indicator of laminar location, which is independent of vertical electrode placement and small changes in electrode placement within the array. The depression of VEP amplitudes during repetitive visual stimulation was accelerated following deprived/contralateral eye stimulation relative to the non-deprived/ipsilateral eye (Second VEP response normalized to first, AVG±SEM; Ipsi 0.59±0.13μV, Contra 0.29±0.02 μV; Repeated measures ANOVA, p < 0.001, F = 47.683, between groups, p = 0.007, F = 15.773, Cohen’s d = 2.31, n = 4 subjects, Fig. 2C). The acceleration of short-term synaptic depression of VEP amplitude suggests that cMD induces an increase in the neurotransmitter release probability at synapses serving the deprived eye, as has been observed in the LGN and visual cortex after brief MD^52, 53^.

### ERs in adult rodent V1

To ask if the sensitivity to αE2 is maintained in the adult visual cortex, we first examined the distribution of canonical ERs. This analysis was performed in adult (>P180), gonadally-intact, male and female LE rats, as gonadectomy has been shown to significantly reduce excitatory synaptic density^11, 12, 54^. Quantitative immunohistochemistry reveals robust expression of ERα and ERβ throughout the brains of adult males and females (Fig. 3A). To ask if ERs in the adult brain are located outside of the nucleus, we compared the distribution of ERα and ERβ to the distribution of DAPI (4’,6-Diamidine- 2’-phenylindole dihydrochloride), a fluorescent stain that binds to AT-rich sequences of DNA. Pearson’s correlation coefficients (PCC) reveal low co-localization between DAPI / ERα and DAPI / ERβ in adult V1 (AVG±SEM; DAPI-ERα: Males 0.048±0.023, Females 0.002±0.037, males vs. females p=0.21, student’s t-test; DAPI-ERβ: Males 0.006±0.024, Females: −0.025±0.037, males vs. females p=0.43, student’s t-test; n=3 males and n=3 females; Fig. 3B). Low co-localization between DAPI and ERα and ERβ was also observed in hippocampal area CA1 of these same subjects, in agreement with previous reports^23, 24^ (AVG±SEM; DAPI-ERα: Males 0.163±0.041, Females 0.223±0.06, males vs. females, p=0.48, student’s t-test; DAPI-ERβ: Males 0.042±0.011, Females: 0.076±0.039; males vs. females p=0.44, student’s t-test, n=3 males and n=3 females; Fig. 3B). Immunolabeling for ERα and ERβ was absent when antibodies were pre-absorbed with antigen (*not shown*), and ERβ staining was absent in the brains of ERβ^−/−^ adult female rats (Representative example of n = 3; Fig. 2A, right^39^). Thus, robust non-nuclear expression of ERs persists throughout the brain of adult male and female LE rats, including primary visual cortex.

**Figure 3.**
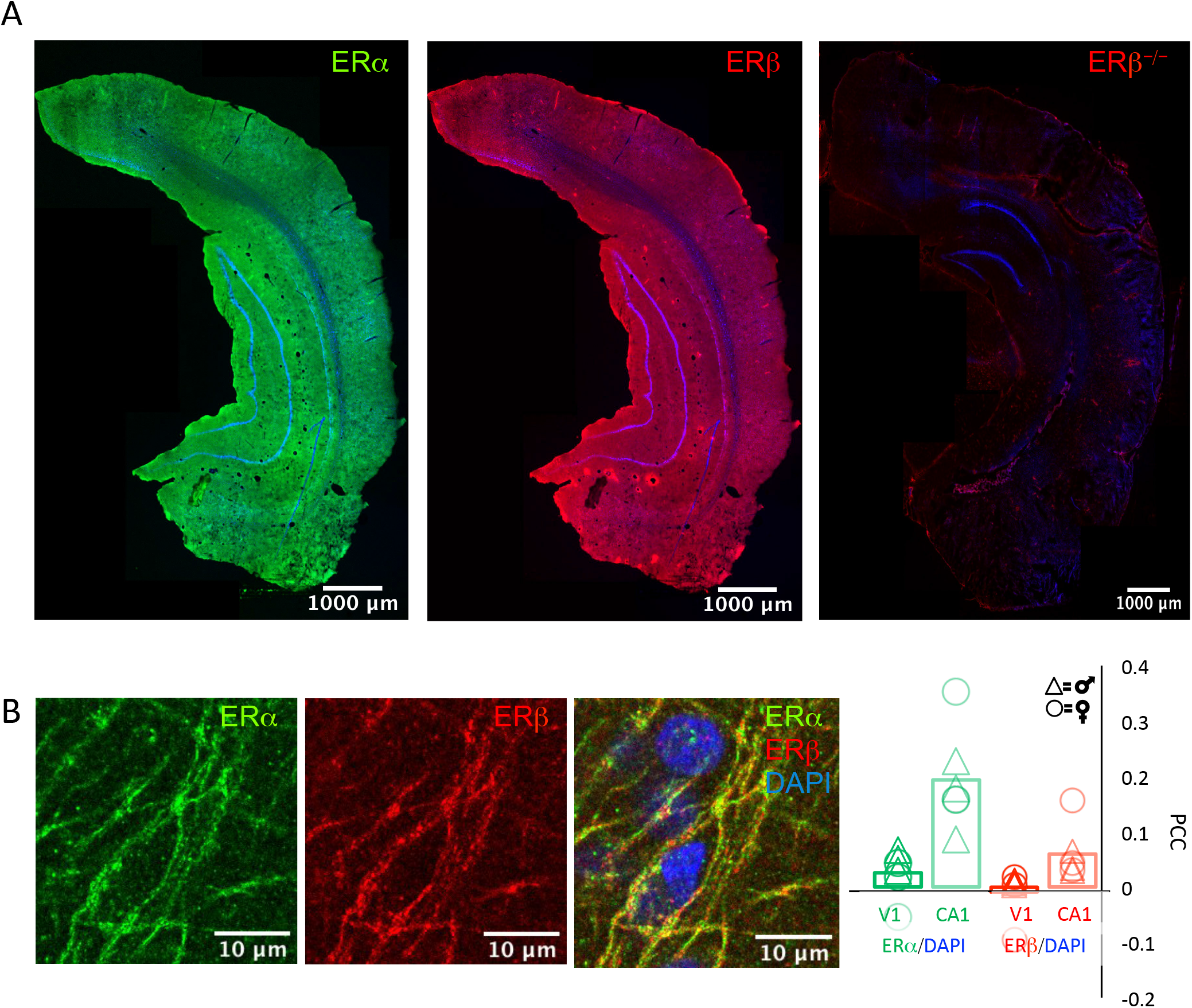
Distribution of estrogen receptors ERα and ERβ in adult male and female rat brains. A. Low magnification single z-section fluorescent micrographs of coronal plane of adult rats. Immunoreactivitity for ERα (left; green) and ERβ (middle; red) in adult male WT and immunoreactivity for ERβ in adult female ERβ^−/−^, counter-stained with DAPI (blue). 5X magnification, ROI 129.17μm × 129.17μm. B. Higher magnification single z-section fluorescent micrographs of triple labeled ERα (green, left) /ERβ (red, middle) and DAPI (blue, right). RO1: V1b 500 μm from surface: 72μm × 82 μm; 40x mag plus 3x digital zoom. Same subjects as in A. C. Pearson’s correlation coefficient (PCC) reveals low correlation between nuclear signal (DAPI) and ERα (green) and ERβ (red) in adult male and female WT rats in V1 (dark green, dark red) and CA1 (light green, light red). Males (triangles) vs. females (circles): student’s t-test: V1 DAPI-ERα p=0.21, DAPI-ERβ: p=0.43, CA1 DAPI-ERα: p=0.48, DAPI-ERβ: p=0.44; n=3 males, 3 females.

The structural and functional deficits induced by prolonged monocular deprivation do not recover spontaneously following removal of the occlusion in adulthood^33, 45^. However, recovery of vision in the amblyopic pathway can be promoted by manipulations that enhance synaptic plasticity in V1b ^55, 56, 35, 57, 58^. To ask if estrogen treatment can enhance structural and functional plasticity in the adult amblyopic V1, we treated with 17α-estradiol (αE2), a stereoisomer of the main circulating estradiol (E2), which induces potent synaptogenesis but is genomically inactive^37^. To confirm the latter, we found that 7 days of αE2 treatment (15 μg/kg, s.c., 1x/day, awake animals) to female and male adults resulted in no change in gonad/body weight (AVG±SEM; female vehicle (veh): 4.10±0.38 vs. female αE2: 5.134±1.09; male veh: 6.01±0.90 vs. male E2: 5.59±0.34, males n=3, females n=3).

### Effect of αE2 on anatomical markers of plasticity

To ask if acute αE2 treatment regulates neuronal excitability in the amblyopic visual cortex, we examined the response to a single dose of αE2 on the expression of the activity-dependent calcium-binding protein parvalbumin (PV), a proxy for the activity of fast-spiking interneurons (FS INs^59^). A dose of αE2, previously shown to induce potent structural plasticity (15 μg/kg, s.c.^37^), was delivered to amblyopic male and female rats. αE2 treatment induced a decrease in the size distribution of PV immunofluorescent puncta in V1b contralateral and ipsilateral to the occluded eye (AVG±SEM; Contra cMD 26.18±2.67 vs. Contra cMD+αE2 24.68±3.61, p<0.001, K-S Test, Cohen’s *d* =0.47; males vs. females, 2-way ANOVA F(1,8) = 4.39, p = 0.07; Ipsi cMD 25.59±1.42 vs. Ipsi cMD+αE2 18.97±3.17; p<0.001, K-S Test, Cohen’s *d*=2.69; males vs. females 2-way ANOVA F(1,8) = 0.41, p = 0.54; n=6, each; Fig.4 A,B). This suggests a reduction in the excitability of FS INs, and is consistent with previous reports that E2 treatment can increase principal neuron excitability and EPSP amplitudes^60, 61, 62^.

**Figure 4:**
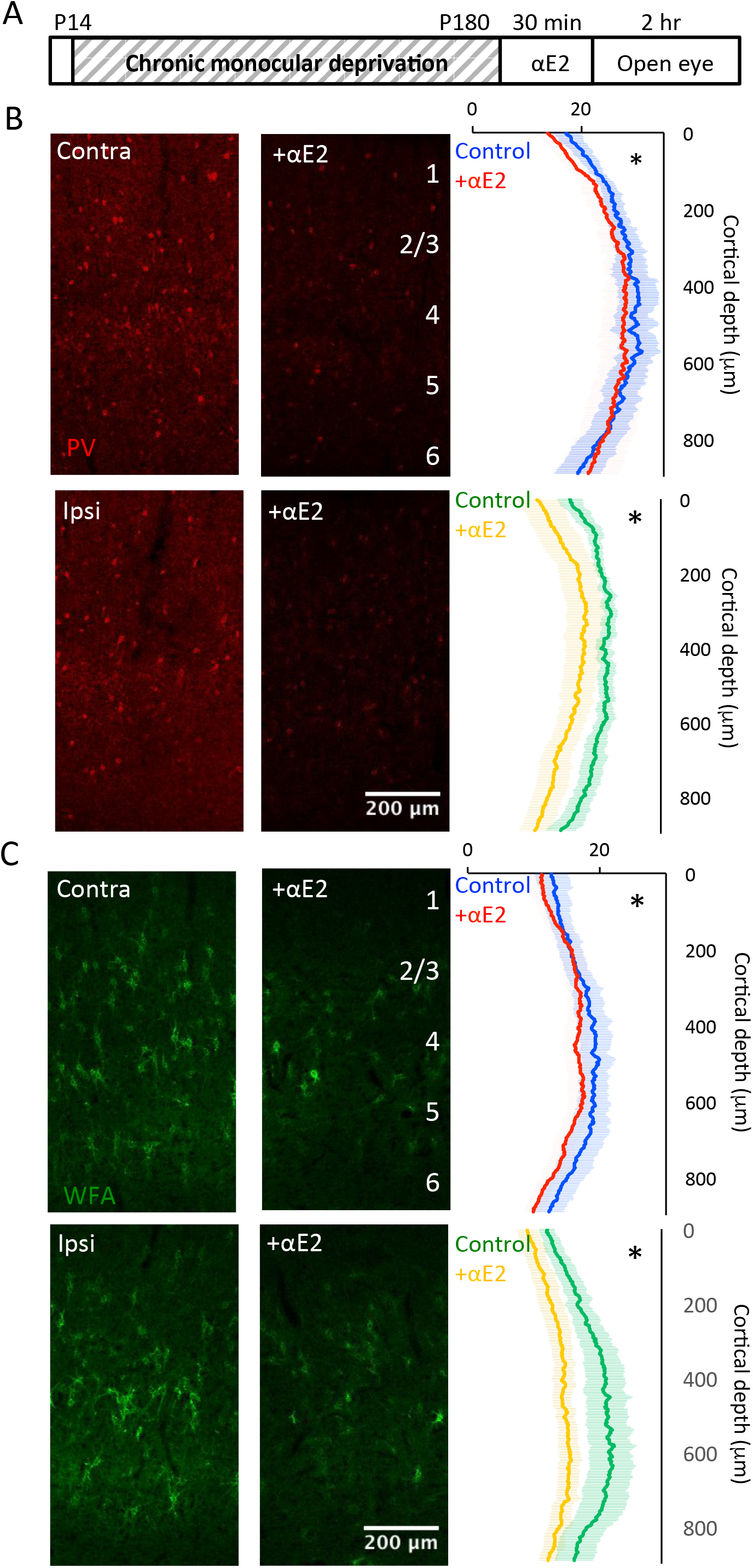
Acute 17α estradiol treatment reduces PV and WFA staining in V1b contralateral and ipsilateral to cMD. A. Experimental timeline. Subjects received monocular deprivation from eye opening (~P14) to adulthood (>P180). 17α estradiol (15 μg/kg, s.c.) was delivered 30 minutes prior to eye opening. B. Left: Fluorescent micrographs of PV distribution in V1b (red; ROI: 900 μm × 500 μm; 10x mag; MIP; 9 × 7.5 μm sections). Right: Average±SEM PV fluorescence intensity in ROI by vertical depth. 17α estradiol significantly decreases PV fluorescence intensity in V1b contralateral and ipsilateral to cMD *p<0.001, K-S Test. Males versus females: 2-way ANOVA Contra: F(1,8) = 4.39, p = 0.07; Ipsi: F(1,8) = 0.41, p = 0.54; n=6 each. C. Left: Fluorescent micrographs of WFA distribution in V1b (green; ROI as in 3b). Right: Average±SEM WFA fluorescence intensity in ROI by vertical depth. 17α estradiol significantly decreases WFA intensity in V1b contralateral and ipsilateral to cMD =p<0.001, K-S Test. Males versus females: 2-way ANOVA Contra: F(1,8) = 3.23, p = 0.11; Ipsi: F(1,8) = 1.69, p=0.23; n=6 each.

Next, we examined the effect of αE2 on the integrity of the extracellular matrix (ECM), the maturation of which is known to contribute to the constraint of structural and functional plasticity with age^63, 64^. Chondroitin sulfate proteoglycans (CSPGs), a primary component of the ECM, are labeled by Wisteria-floribunda agglutinin (WFA) binding to N-acetyl-D-galactosamine in the chondroitin sulfate chain. αE2 treatment of cMD subjects (15 μg/kg, s.c.) induced a decrease in the distribution of fluorescein-WFA staining in V1b contralateral and ipsilateral to the occluded eye (AVG±SEM; Contra cMD 16.63±2.08 vs. Contra αE2 14.85±1.82, p<0.001 K-S Test, Cohen’s *d* = 0.91; males versus females: F(1,8) = 3.23, p = 0.11, 2-way ANOVA; Ipsi cMD 18.23±2.79 vs. Ipsi αE2 12.62±1.56, p<0.001 K-S Test, Cohen’s *d* = 2.48; males versus females: F(1,8) = 1.69, p=0.23, 2-way ANOVA; n=6, Fig.4 C,D). Thus, αE2 treatment induces changes in the adult amblyopic cortex that are predicted to promote synaptic plasticity.

### Quantitative immunofluorescence of PSD95 and pS831 following αE2

To ask if αE2 treatment impacts the density of excitatory synapses, we again examined the intensity and distribution of PSD95 labeling. αE2 treatment of cMD subjects (15 μg/kg, s.c., awake subjects) induced a significant increase in the size distribution of PSD95 immunoreactive puncta in V1b contralateral, but not ipsilateral, to the occluded eye (Contra cMD 0.087±0.01 μm^2^ vs. Contra αE2 0.088±0.005μm^2^, p<0.001 K-S Test, Cohen’s *d* = 0.13; males versus females: F(1,2) = 0.98, p = 0.34, 2-way ANOVA; Ipsi cMD 0.088±0.01 μm^2^ vs. Ipsi αE2 0.09±0.001 μm^2^; males vs. females, F(1,12) = 0.11, p = 0.75, 2-way ANOVA), but no change in PSD95 number (AVG±SEM; Contra cMD 320.65±54.46 vs. Contra αE2 364.03±47.96; males vs. females, F(1,12) = 1.84, p = 0.95, 2-way ANOVA; Ipsi cMD 297.87±53.71 vs. Ipsi αE2 389.46±64.07; males vs. females, F(1,12) = 4.27, p = 0.06, 2-way ANOVA; n=8 each, Fig.5B). In contrast, we observed a decrease in the size distribution of pS831 immunoreactive puncta in both hemispheres (AVG±SEM; Contra cMD 0.078±0.006 μm^2^ vs. Contra αE2 0.071±0.004 μm^2^, p<0.001 K-S Test, Cohen’s *d* = 1.37; males versus females: F(1,12) = 0.138, p=0.72, 2-way ANOVA; Ipsi cMD 0.077±0.006 μm^2^vs. Ipsi αE2 0.076±0.005 μm^2^, p<0.001, K-S Test, Cohen’s *d* = 0.18; males versus females: F(1,12) = 0.01, p=0.91, 2-way ANOVA) but no change in puncta number following αE2 treatment (Contra cMD 283.68±64.10 vs. Contra αE2 359.06±69.38; males versus females: F(1,12) = 1.38, p=0.26, 2-way ANOVA; Ipsi cMD 305.01±66.20 vs. Ipsi αE2 358.53±70.82; males versus females: F(1,12) = 2.16, p=0.17, 2-way ANOVA; n = 8, Fig.5C). This suggests that αE2 treatment increases the size of existing excitatory synapses in the adult V1 via a pathway that is independent of CaMKII/PKC phosphorylation of GluA1.

**Figure 5.**
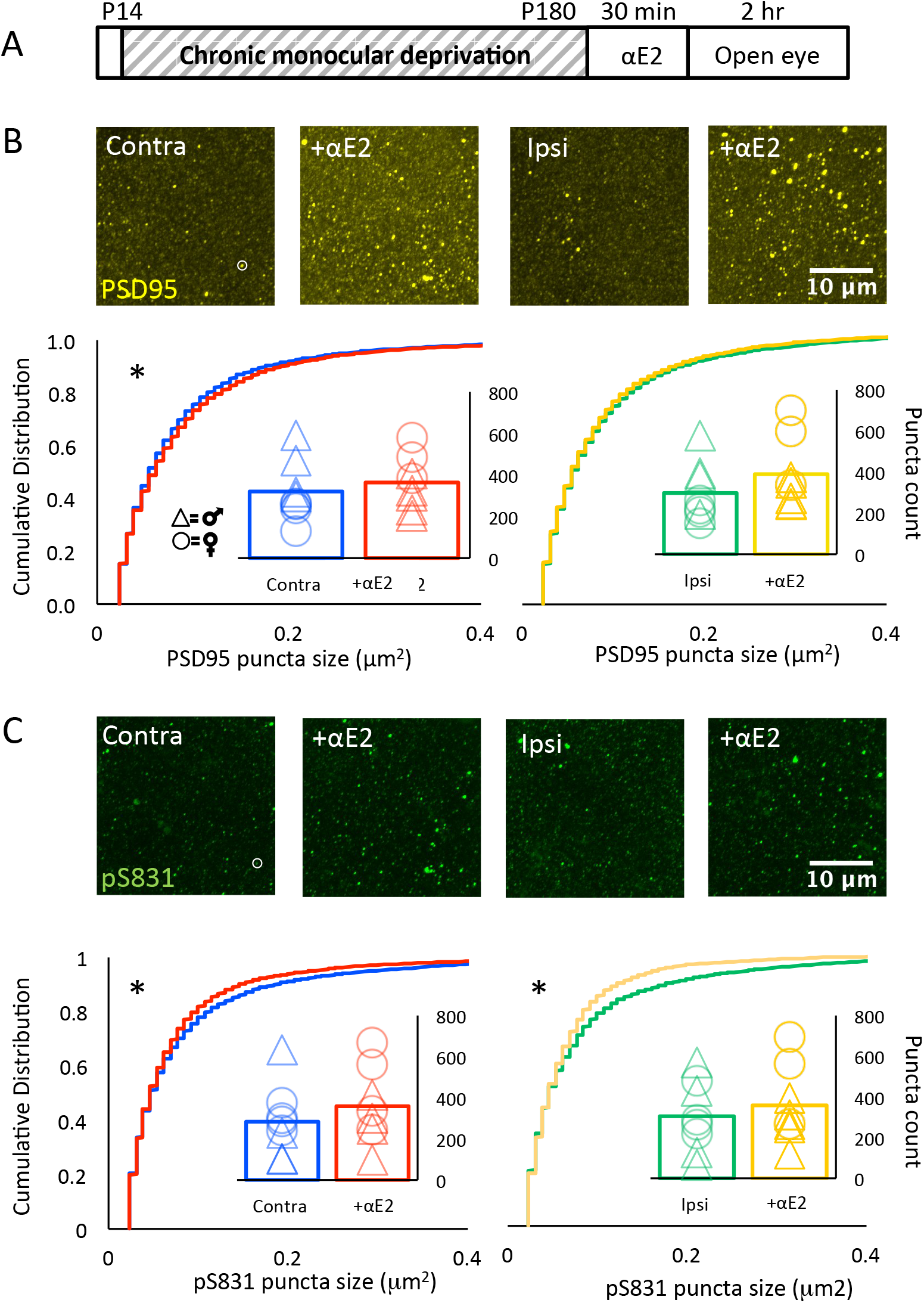
Acute 17α estradiol treatment increases PSD95 and decreases pSer831 puncta size. A. Experimental timeline. Subjects received monocular deprivation from eye opening (~P14) to adulthood (>P180). 17α estradiol (15 μg/kg, s.c.) was delivered 30 minutes prior to eye opening. B. Top: Fluorescent micrographs of PSD95 immunoreactivity (yellow, representative puncta in white circle) in V1b±17α estradiol contralateral (left) and ipsilateral (right) to cMD (V1b; 500 μm from surface; ROI: 28.34 μm × 28.34 μm × 40 μm, 100x mag with 3x digital zoom; MIP; 40 × 1μm z-steps). Bottom: Cumulative distribution reveals a significant increase in PSD95 immunoreactive puncta size in V1b contralateral (left), but not ipsilateral (right) to the deprived eye, *=p<0.001, K-S Test. Males (triangles) vs. females (circles): 2-way ANOVA Contra: F(1,12) = 0.98, p = 0.34, Ipsi: F(1,12) = 0.11, p = 0.75. No change in PSD 95 immunoreactive puncta number (insets). Males (triangles) vs. females (circles): 2-way ANOVA Contra: F(1,12) = 1.84, p = 0.95, Ipsi: F(1,12) = 4.27, p = 0.06, n=8. C. Top: Fluorescent micrographs of pS831 immunoreactivity (green, representative puncta in white circle) in V1b±17α estradiol contralateral (left) and ipsilateral (right) to cMD. ROI as in 4b. Bottom: Cumulative distribution reveals a significant decrease in pS831 immunoreactive puncta size in V1b contralateral (left) and ipsilateral (right) to the deprived eye, *=p<0.001, K-S Test. Males (triangles) vs. females (circles): 2-way ANOVA Contra: F(1,12) = 0.138, p=0.72; Ipsi: F(1,12) = 0.01, p=0.91. No change in pS831 immunoreactive puncta number (insets). Males (triangles) vs. females (circles): 2-way ANOVA Contra: F(1,12) = 1.38, p=0.26, Ipsi: F(1,12) = 2.16, p=0.17, n=8.

### Effect of αE2 on experience-dependent synaptic plasticity in cMD subjects

To ask if αE2 treatment enhanced functional plasticity in adult amblyopic V1, we adapted a visual stimulation protocol known to induce stimulus-selective response potentiation (SRP) of visual responses in binocular mice^65^. Repetitive presentation of high contrast gratings of a single orientation (200-500 phase reversals a day over 5-7 days) induces a two-fold increase in the amplitude of the VEP recorded in layer 4 in awake mice. The visual response potentiation is highly selective for the characteristic of the visual stimulus including orientation and temporal frequency, thereby reproducing the specificity of many forms of visual perceptual learning^66^.. In addition our previous work demonstrates that a truncated stimulation protocol, in which subjects receive 100-200 phase reversals of a high contrast grating, is subthreshold for the induction of stimulus-selective response potentiation (SRP) in anesthetized rats^66^. However, this truncated stimulation protocol is sufficient to induce SRP in amblyopes following manipulations that increase plasticity in the visual cortex. To ask if αE2 treatment regulates the response to this protocol, we presented repetitive visual stimulation binocularly (200 phase reversal of 0.05 cycles per degree (cpd), 100% contrast gratings, 45 degrees, reversing at 1 Hz). After 24 hours, we compared monocular VEP amplitudes in response to the familiar and novel grating orientations (45 and 135 degrees respectively). As expected, we observed no increase the amplitude of the VEP acquired from the previously deprived or non-deprived eye in vehicle-treated controls (AVG±SEM; non-deprived, familiar: 22.37±1.28 μV, novel: 22.71±2.55 μV; males versus females: F(1,4) = 1.09, p = 0.36, 2- way ANOVA; deprived, familiar: 21.65±3.59 μV, novel: 19.99±3.36 μV; males versus females: F(1, 4) = 1.19, p = 0.34, 2-way ANOVA; n=4, Fig. 6B,C). However, αE2 treatment prior to the initial visual stimulation enabled a significant increase in VEP amplitude in response to stimulation of the non-deprived eye (AVG±SEM; familiar: 36.10±6.38 μV, novel: 27.11±4.93 μV; *p = 0.024 two- tailed paired t-test, Cohen’s *d* = 0.13; males versus females: F(1,8) = 2.05, p = 0.19, 2-way ANOVA; n=6 subjects Fig. 6B). Importantly, the increase in the amplitude of the non-deprived eye VEP was stimulus- selective, as the increase was observed in response to familiar, but not novel, visual stimulus orientations (Fig. 6B). In contrast, no change was observed in the amplitude of the VEP acquired from the previously deprived eye (AVG±SEM; familiar: 25.56±4.49 μV, novel: 29.154±3.80 μV; males vs. females, F(1, 8) = 1.59, p = 0.24, 2-way ANOVA; n=6, Fig. 6C). Together this demonstrates that adult amblyopic V1 retains sensitivity to E2, and αE2 treatment specifically enhances plasticity at the spared synapses serving the non-deprived eye.

**Figure 6.**
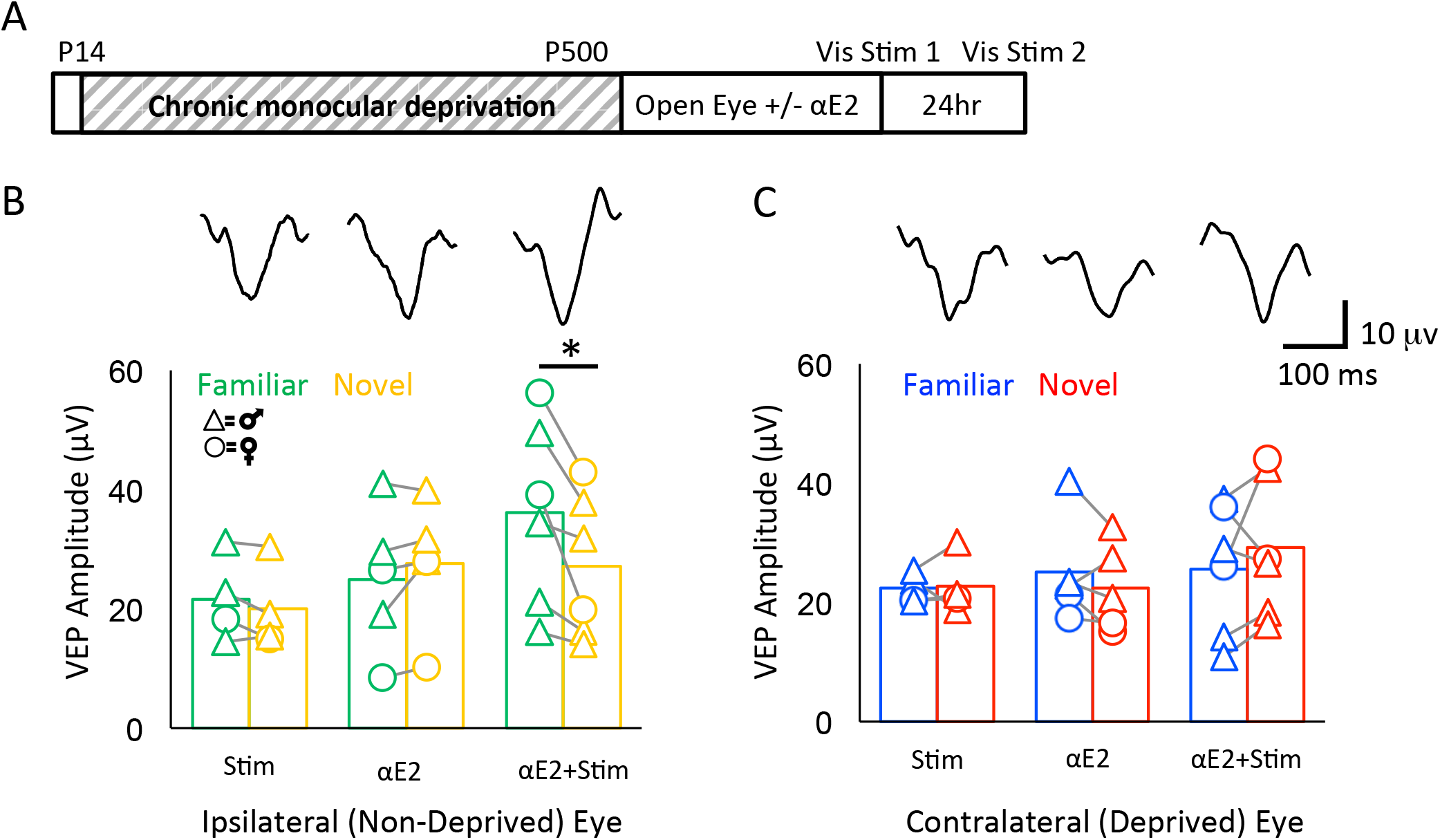
Acute 17α estradiol promotes stimulus-selective response potentiation (SRP) of the non-deprived eye VEP. A. Experimental timeline. Subjects received monocular deprivation from eye opening (~P14) to adulthood (>P180). 17α estradiol (15 μg/kg, s.c.) was delivered 30 minutes prior to eye opening and initial binocular visual stimulation (200 phase reversals at 1 Hz of 0.05 cycle per degree 100% contrast gratings at 45 degrees). VEPs were acquired from layer 4 in response to familiar (45 degrees) and novel (135 degrees) grating orientations 24 hours after the initial visual stimulus. B. Average VEP amplitude evoked in ipsilateral (non-deprived) eye in response to familiar (green) and novel (yellow) visual stimuli. No difference in response to familiar versus novel stimuli in subjects that received visual stimulation or αE2 alone. Significant increase in response to familiar versus novel visual stimuli in subjects that received αE2 prior to initial visual stimulation. Inset: representative VEP waveforms in response to the familiar stimulus in vis stim only (left) αE2 (center) and αE2 + vis stim (right). *p<0.05, two-tailed paired t-test. Stim n=4, αE2 n=5, αE2+Stim n=6. Males (triangles) vs. females (circles): 2-way ANOVA F(1,8) = 2.05, p = 0.19. No change in average VEP amplitude evoked in contralateral (deprived) eye in response to familiar (blue) and novel (red) stimuli in any condition. Inset: Representative VEP waveforms in response to the familiar stimulus in vis stim only (left) αE2 (center) and αE2 + vis stim (right). Stim n=4, αE2 n=5, αE2+Stim n=6. Males (triangles) vs. females (circles): 2-way ANOVA F(1,8) = 1.59, p = 0.24).

## Discussion

The enhancement of structural and functional plasticity by E2 is well-documented in adult hippocampus, hypothalamus and frontal cortex^2, 7, 11, 67, 68^. Here we show that estradiol treatment can promote plasticity in amblyopic visual cortex by quantifying the effects of αE2 on the structure and function of the binocular region of the adult visual cortex following chronic monocular deprivation. A single dose of αE2 reduced the expression of PV, reduced the integrity of the ECM, and increased the expression of the postsynaptic scaffold PSD95. Furthermore, αE2 treatment enhanced experience-dependent plasticity in the amblyopic cortex, promoting stimulus-selective response potentiation in the pathway served by the non-deprived eye. Our results demonstrate that the adult amblyopic cortex retains responsiveness to E2, and suggests that the sensitivity to acute αE2 may be determined by synaptic properties that reflect the history of activity at the synapse.

The initiation of MD initiated at eye opening (cMD) mimics the severe deprivation amblyopia caused by the present of a unilateral congenital cataract^33^. cMD induces a severe asymmetry in the structure and function of the cortical circuitry serving the deprived eye versus non-deprived eye ^33, 35, 66^ and therefore contrasts with MD initiated during the critical period which induces a more modest structural and functional changes. The cMD model allowed us to compare the response to acute αE2 treatment in the compromised pathway serving the deprived eye and the intact pathway serving the non-deprived eye. Intraocular delivery of a trans-neuronal tracer following cMD revealed the expected decrease in thalamocortical inputs serving the deprived eye, reflected in the deprivation-induced reduction in the strength of thalamic input to the cortex^45^. Nonetheless, normal PSD95 and pS831 labeling was observed after cMD, suggesting an increase in other classes of excitatory synapses following the loss of thalamocortical inputs. Indeed, MD during the critical period induces a rapid depression of deprived eye responses followed by a slowly emerging enhancement of non-deprived eye responses^46^. The expansion of thalamocortical input into V1m may also contribute to the increase in non-deprived eye response strength.

The therapeutic potential of E2 /αE2 in the adult brain critically depends on the distribution and concentration of ERs. The persistence of robust, non-nuclear ER expression after menopause/estropause is well documented in primate and rodent hippocampus, hypothalamus, and frontal cortex. However, there has been little consensus on the distribution or role of ERs in primary sensory cortices of adults^29, 30, 31, 32, 48^. Our results demonstrate that robust ERα and ERβ expression persists V1 of adult male and female LE rats. The majority of ERα and ERβ labeling did not co-localize with a nuclear marker in the primary visual cortex or hippocampus, the latter consistent with previous reports of high non-nuclear receptor expression in CA1 of adult female rats^23, 24^.

In addition to the reduction in circulating sex hormones, the maturation of extracellular matrix (ECM) is known to constrain structural and functional synaptic plasticity in adult circuits^42, 64^. A single dose of αE2 reduced the integrity of the ECM throughout the visual cortex of amblyopic adults, thereby mimicking other interventions that enhance plasticity in adult V1^42, 57, 63, 70^ αE2 also reduced the expression of PV, a proxy for the excitability of FS INs^13^. However, the reduction in ECM integrity and FS IN excitability did not induce a global enhancement of plasticity in synapses in V1.

Acute E2 delivered *in vivo* or *ex vivo* induces robust spinogenesis in the hippocampus of young adult male and female rats^11, 25, 69, 71^ and this effect may be lost with age or following OVX^72^. Brief E2 treatment of hippocampal slices from young males and middle-aged OVX females induces an increase in F-actin, without a change in PSD95 puncta number^14, 73^. Similarly, Golgi stains of mouse hippocampus from OVX females (P42) reveal an increase in the number of large, mushroom-type spines following repetitive E2 treatment (1x day / 5 days^74^). Following acute delivery of αE2, we observed an increase PSD95 puncta size in the adult visual cortex, but no change in the number of PSD95 puncta. Together, this suggests that the estradiols induce the genesis of new excitatory synapses in young adults, and the growth/expansion of pre-existing excitatory synapses in older brains. Importantly, we observed no difference in PSD95 puncta size or number in males versus females, and low variability within the female cohort. This suggests minimal impact of the estrous cycle phase on PSD95 expression in our adult rats.

αE2 treatment induced an increase in the size of PSD95 puncta, suggesting that αE2 treatment alone may induce a modest strengthening of excitatory synapses, as has been shown in response to E2^76^. Additionally, acute E2 lowers the threshold and increases the magnitude of LTP induced by theta burst stimulation in the hippocampus of OVX rats^67^. However, the absence of an increase in pS831 following αE2 treatment demonstrates that the increase in the size of excitatory synapses occurs independently of CaMKII/PKC signaling. Together this suggests that estradiol treatment engages a pathway parallel to that engaged by LTP, and is consistent with the observation that the response to acute E2 in hippocampus, including polymerization of actin in dendritic spines and an increase in excitatory synaptic strength, promotes rather than occludes subsequent LTP. An E2-induced increase in excitatory synaptic strength may also underlie reports of in increased LTD magnitude^25, 76^.

E2 treatment increases the amplitude and frequency of mEPSCs in rat hippocampal CA1 pyramidal neurons in both sexes, indicative of regulation of pre- and postsynaptic function respectively^43^. Importantly, E2-induced changes in mEPSC frequency are limited to synapses with an initially low probability of neurotransmitter release, suggesting that E2 enhances synaptic function by increasing presynaptic release probability^15^. A similar synapse-specific effect of αE2 was observed in the adult amblyopic visual cortex, with enhancement of activity- dependent plasticity limited to the pathway serving the non-deprived eye. The observation that the VEPs acquired from the deprived eye depress more rapidly than the non-deprived eye suggests that chronic monocular deprivation may increase the probability of neurotransmitter release at deprived-eye synapses, as observed following brief MD in the thalamus and cortex^52, 53^. The increase in release probability of neurotransmitter at synapses serving the deprived eye may occlude the enhancement of plasticity by αE2 and underlie the selectivity for spared inputs in the amblyopic cortex. The robust expression of ERs in the adult visual cortex and the enhancement of plasticity of synapses in the intact, non-deprived eye pathway by αE2 suggests that estradiol treatment could be employed to promote the plasticity of spared inputs around a scotoma or a cortical infarct. The selective enhancement of plasticity at synapses with initially low release probability also suggests the possibility that sensitivity to αE2 could be acutely manipulated following induction of activity-dependent changes in the probability of neurotransmitter release.

## Author Contribution Statement

DCS performed the experiments in Figures 1, 2A, 2B, 3, 4, and 5. CLL performed the experiments in Figures 2C and 6. MAKR provided the ERβ^−/−^ rats and contributed to data interpretation. DCS, CLL and EMQ designed the experiments and wrote the manuscript.

## Competing Interests Statement

Deepali C. Sengupta declares she has no competing interests. Crystal L. Lantz declares she has no competing interests. M.A. Karim Rumi declares he has no competing interests. Elizabeth M. Quinlan declares she has no competing interests.

## Ethical approval Statement (duplicated in methods section)

All procedures were approved by the University of Maryland Institutional Animal Care and Use Committee and were carried out in accordance with the Guide for the Care and Use of Laboratory Animals.

## Data Availability Statement

The datasets generated during the current study are available from the corresponding author on reasonable request.

## Acknowledgements

This work was supported by NIH R01EY016431 (to EMQ), and an ARCS-MWC Foundation Fellowship, a UM NACS Hodos Dissertation Completion Fellowship and a Cosmos Club Foundation Scholars Fellowship (to DCS).

